# Can constraint closure provide a generalized understanding of community dynamics in ecosystems?

**DOI:** 10.1101/2020.01.28.924001

**Authors:** Steven L. Peck, Andrew Heiss

## Abstract

Since the inception of the discipline, understanding causal complexity in ecological communities has been a challenge. Here we draw insights from recent work on constraint closure that suggests ways of grappling with ecological complexity that yield generalizable theoretical insights. Using a set of evolutionary constraints on species flow through ecological communities, which include: selection, species drift, dispersal, and speciation, combined with multispecies interactions such as mutualistic interactions, and abiotic constraints, we demonstrate how constraint closure allows communities to emerge as semi-autonomous structures. Here we develop an agent-based model to explore how evolutionary constraints provide stability to ecological communities. The model is written in Netlogo, an agent based-modeling system, with advanced tools for manipulating spatially structured models and tools for tracking pattern formation. We articulate ways that ecological pattern formation, viewed through the lens of constraint closure, informs questions about stability and turnover in community ecology. The role of the chosen constraints was clear from the simulation results. It took the shape of both inducing stability and creating conditions for a more dynamic community with increases in species turnover through time. Key ecological and evolutionary variables showed overall stability in the landscape structure when plotted against the number of constraints, suggesting that these evolutionary forces act as constraints to the flow of species in such a way that constraint closure is achieved effecting semi-autonomy.

**Author Summary:** Ecosystems are among the most complex structures studied. They comprise elements that seem both stable and contingent. The stability of these systems depends on interactions among their evolutionary history, including the accidents of organisms moving through the landscape and microhabitats of the earth, and the biotic and abiotic conditions in which they occur. When ecosystems are stable, how is that achieved? Here we look at ecosystem stability through a computer simulation model that suggests that it may depend on what constrains the system and how those constraints are structured. Specifically, if the constraints found in an ecological community form a closed loop, that allows particular kinds of feedback may give structure to the ecosystem processes for a period of time. In this simulation model, we look at how evolutionary forces act in such a way these closed constraint loops may form. This may explain some kinds of ecosystem stability. This work will also be valuable to ecological theorists in understanding general ideas of stability in such systems.

## Introduction

Ecological theorists have generated several yet unresolved disputes that try to untangle the difficulty of understanding the nature of complex ecological communities–including the question of whether the idea of a community in such systems is coherent. For example, questions about ecological community structure and how it is maintained have been argued about for over a hundred years. These questions are still taking their current principle divisions from camps first staked out by Clements [1] and Gleason [2] in the early decades of the 20th Century: Are communities assembled non-randomly, conditioned on certain possibilities of co-occurrence and structured by general ecological regularities, or are they more randomly constructed entities brought together by a complex combination of abiotic factors, available species pools, competition, and other ecological processes that suggest a strong element of contingency [3, 4]?

Currently, there does not seem to be a unified body of theory and practice that explains most features of community structure, and many of the theories such as those above are handled separately. The hunt for generality has also been fraught (See Elliott-Graves [5] for a detailed overview). Theories in the discipline often seem a piecemeal confederation of contingent ideas that carve nature mid-bone, as it were, often focused on specific natural systems that provide only limited empirical and theoretical support among other differently structured ecological communities. It is clear that the theoretical constructions used by ecologists have been crucially instrumental in the development of research programs and have allowed progress in understanding community ecology. However, even basic concepts in the discipline are often viewed differently by different research groups that are often divided into different camps, often to the point that there are even debates about the usefulness of the concept of community ecology. For example, the following questions are still being debated. Are they individuals [6–15]? Which ecological theories best describe the dynamics of their structure, maintenance, and development [9, 10, 12, 14, 15]? Do they have boundaries [6, 16]? Are they stable [17]? How are communities assembled, and are they so contingent that each community must be examined as an individual case study [18–24]? Are ecosystems even real entities [25, 26]?

These questions are all interrelated, yet the abundant theories used to tame and understand these ecological systems rarely provide a unified or coherent picture among theories. Moreover, in order to make progress in the idea of an ecological community, it seems essential to discover generalities that hold across the numerous possible biotic configurations that comprise these individual community assemblages. It is also important to view this project from multiple perspectives in order to tackle the inherent complexity and provide a unified or coherent picture among theories for a more unified conception of community ecology. There have been many attempts to do so, and the importance of these efforts has been recognized, [10, 12, 15, 21, 27–32], though conditioned on some skepticism about the possibility of finding well-supported law-like regularities in community ecology [20, 33–35].

### The Piecemeal Nature of Ecological Theories

Almost from the inception of ecology as a discipline, piecemeal theories emerge that propose ways to handle its untamed complexity. Table 1 provides a sampling of some of these theories. Note that the structure of these theories is such that each takes a particular, narrowly focused set of ecosystem patterns and processes from which it attempts to reduce complexity by simplifying the target system to a few aspects extracted from the whole. Scheiner and Willig [36] provide an excellent overview of how these theories function in a hierachieal structure that moves from general theories of the discipline of ecology, to constitutive theories such as the example frameworks found in Table 1, to quantitive models of specific processes that target specific aspects of these theories.

**Table 1.** Sampling of Ecological Theories.

|  |
| --- |
| Network Theory [37,38] |
| Food webs [39,40] |
| Metapopulation Theory [?,41] |
| Niches and Niche Construction [42–44] |
| Island Biogeography [?,45] |
| Hubbell’s Unified Neutral Theory<br>of Biodiversity and Biogeography [46,47] |

For example, food web theory looks at interactions among species that feed in trophic hierarchies; whereas metapopulation theory considers movement among organisms among small populations scattered in a matrix of uninhabitable or poor quality habitat; and neutral theory examines community construction from the perspective of available species pools exploiting contingent opportunities to commandeer openings in resource availability. As Scheiner and Willig list in their paper, many other such ecological theories could have been included in Table 1. However, in what follows we will focus on community ecology, a constitutive theory within the general theory of ecology. This focuses on the interrelationships of biotic and abiotic factors of locally interacting organisms and their contexts.

### Community Ecology

Community ecology seems to lack a grounding generalization that would inspire more confidence that researchers have articulated a well defined domain and framework for community ecology. Most draw only lightly on defining the domain over which they hold sway and are composed of a patchwork of other constitutive theories.

Pragmatically, it should be noted that these constructions work well enough to make these frameworks useful in providing grounding for experimental design and data analyses to support making progress in community ecology. This may be why community ecology has seemed so elusive. The piecemeal nature of its theories, the lack of formal attributes that allow a clear and complete articulation of its domain, and other aspects that tie it to ecological theory, have all made the discipline poorly supported theoretically compared to other frameworks in ecology. Such frameworks do much of the work of theories; they organize thought by providing a structure to inform experimental design, data gathering, and analysis. They also provide predictions that set up expectations and interpretations by using various kinds of baseline models.

However, integrating these frameworks, given the complexity of ecosystems, has been problematic. In part, because we want a generalized structure of an ecological community to contain all of these: food webs, niches, metapopulations, contingent openings in niche resources, and the other framework-based articulations of ecosystem components and processes. All of these frameworks have been successful in providing information about certain aspects of community structure. They provide insights into the nature of these systems that allow them to be better understood, analyzed, and predictions made based on the assumptions and relationships captured in the constituent models. However, an overarching and general articulation of ecosystem communities has been hard to come by because of its inherent complexity and empirical intractability. This is important because although we want to acknowledge that the models and theories of community ecology have done much to open our understanding to some of the aspects of the science, to understand ecosystems as ecosystems, we are going to have to include their complexity qua complexity if they are to be understood holistically rather than in piecemeal fashion [48].

In short, we are confronted with the problem of studying highly complex systems in which regularities are tamed by conceptual frameworks that, while doing real work in establishing a structure from which to view ecosystem processes, work piecemeal and do not generalize easily among differing frameworks. This is not intended as a criticism, and we recognize that it may be how ecological regularities are best handled, given the limitations of fieldwork. Nevertheless, a more general theory of community ecology would be desirable.

In this paper, we combine two recent approaches that used together suggest a promising way to think more about community ecology and offer the potential for a more generalized theory of community ecology. This work fits well with Odenbaugh’s articulation of an integrating theory [49] in which a simulation model is used to support our articulation of this theory. First, we consider the proposal by Vellend (2016) [15] to focus on a small set of processes that provide a basis for ecosystem theory, similar to the way that theoretical population genetics has been fashioned to focus on a small set of processes. Second, we explore the role of functional constraints in defining autonomous and semi-autonomous systems, of which structured ecological communities may, arguably, be considered [8, 50–54].

### Generalizing Ecological Community Theory

The role of evolution in ecological communities is considered fundamental to the processes of community formation. As the philosopher of science, Jay Odenbaugh, points out, two formulations can be recognized [49], “*current* ecological processes are in play because of *past* ecological processes” and “*current* evolutionary processes are in play because of *past* ecological processes” [Italics in the original]. Mark Vellend proposes we look at ecological communities in light of four evolutionary processes found in a wide variety of ecosystems: species selection, species drift, dispersal, and speciation [15, 55]. These were chosen to mirror the processes found in basic population genetics: selection, genetic drift, dispersal and mutation. These community-level processes focusing on species are thought to capture the means whereby most communities change. Vellend argues that population genetics has made significant advances in understanding how genetic concerns play out in populations by focusing on simple models of these activities. He suggests that by focusing on these fundamental aspects population genetics, has been able to make significant theoretical inroads in understanding how genes in populations are structured and behave. Can community ecology likewise benefit by focusing on a similar set of ecosystem characteristics? Vellend (2016) [15] suggests that this subset of possible processes covers much of the conceptual territory being explored in community ecology.

We suggest that his insights might be expanded by considering communities as autonomous structures conditioned on the constraints imposed by the processes he suggests (species selection, species drift, dispersal, and speciation); by adding three additional forces: mutualism among a subset of species, abiotic boundaries, and patch structure and dynamics. This set seems to capture a broad and well-studied set of constraints in ecosystem communities. Focusing on constraints may be an especially useful approach for ecological generalization as it provides ways to theoretically consider general outcome characteristics like stability, expected species turnover rates, and other features of community dynamics like those noted by Gleason and Clements about how communities are assembled and maintained. How constraints allow the emergence of autonomous systems, which we turn to next, will suggest how these ideas can work together to inform ecology.

### Autonomous Systems

One way to approach a general idea of ecosystem functioning can be drawn from Moreno and Mossio’s (2015) [53] work on biologically autonomous systems. Even through the question of whether ecological communities are autonomous entities is unsettled, they identify three dimensions of general biological autonomy.

The first dimension of biological autonomy is the capacity for self-determination, meaning that the grounding, normativity, teleology, and functionality come from the system itself. For this to occur, a closure of constituent constraints is required. Fig 1 illustrates a minimal set of requirements for the occurrence of constraint closure such that components modulate the flow of energy and materials within the system. These provide a sustainable feedback loop that allows the system to achieve coherence for a period of time, *τ*_1_.

**Fig 1.**
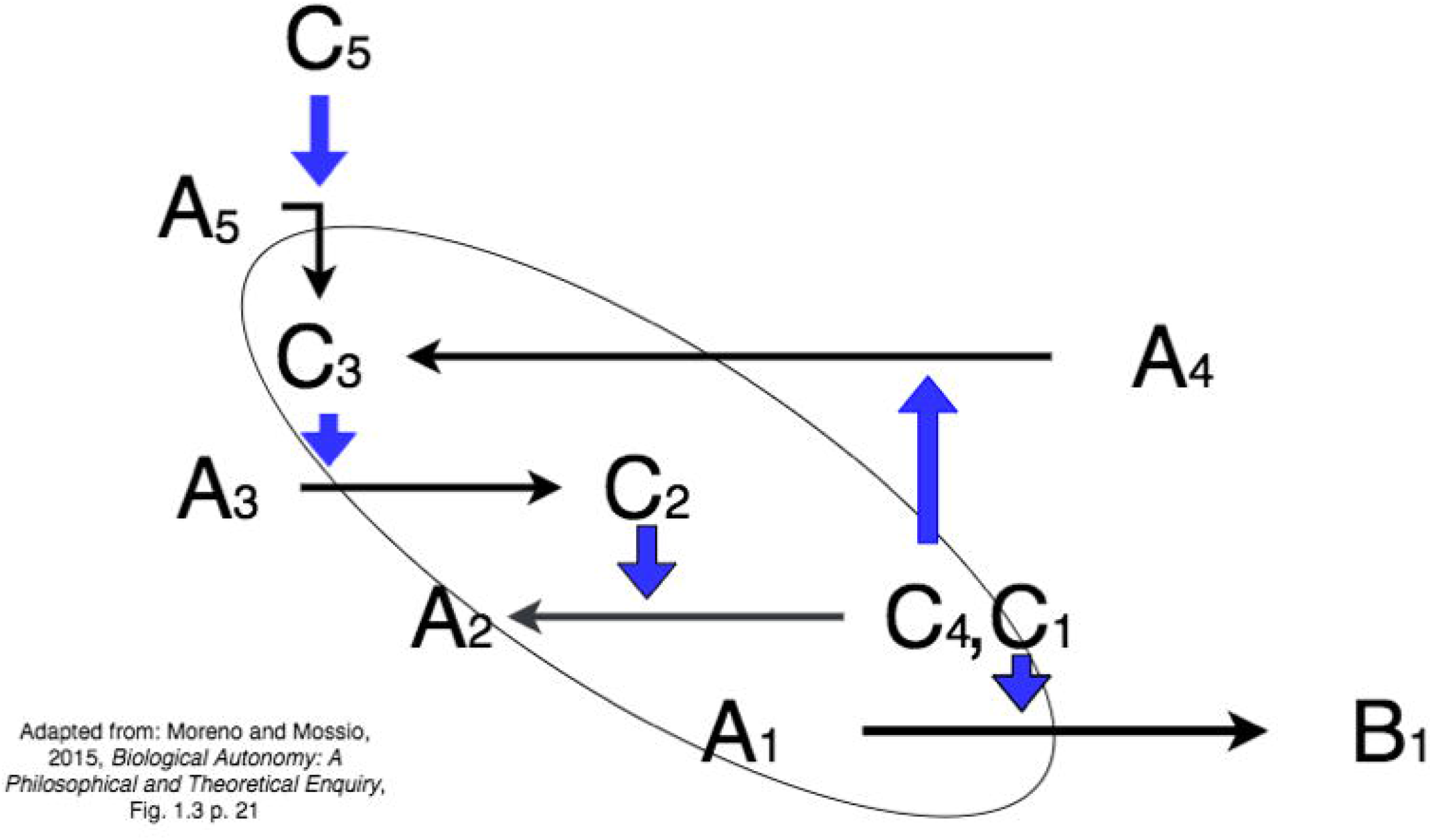
Constraint Closure. Constraint closure is described as constraints on the flow of matter or energy that form a closed loop. This is a necessary step in the formation of an autonomous system. Such a configuration of constraints provides a sustainable feedback loop that allows the system to achieve coherence for a period of time, *τ*_1_. Redrawn from Figure 1.3, p. 21, Moreno and Mossio’s (2015) [53]

The second dimension is found in the interactions with processes conditioned on the larger environment that provides the energy and materials used by the system. For example, in an ecological community, these interactions can take the form of disturbances that might restructure the system through things like direct human influence, invasive species, diseases, or abiotic influence such as fire, weather-related events, and other non-biological influences.

The third dimension of biological autonomy includes the historical aspects of the system. For example, in an ecosystem it might include past arrival of new species, ecological processes, evolutionarily forces, contingent factors, co-evolutionary adaptations, and other biotic and abiotic influences.

Given the above, there are reasons to think that ecological communities might be autonomous systems. Energy flow within the system is maintained and channeled within the system itself by its components. For example, long-term community stability has been shown to be structured by a number of well-studied ecosystem processes, e.g., competitive exclusion of species restructuring particular niches, networks of mutualistic relations, typical ecological interactions like predator-prey relations, etc.

Likewise, there are also reasons to think they are not. Do the contingent aspects of ecologies affect the possibility of their being autonomous? Many of the structures we see in ecosystems appear to be arbitrary members of a set derived from a contingent species pool, what is autonomous in such a system? Is it, perhaps, the fungible functional role that particular species play in an ecosystem and that subsequently structure the patterns and processes we observe? Or does it depend on long-term coevolutionary relationships among particular species? Both may be important, and therefore, some generalities in the form of regularities that span different kinds of ecosystems may qualify these as autonomous systems at least for some interval *τ*. One particular problem is assigning boundaries to ecological communities. This seems to be an important unexplored step in exploring these systems as autonomous systems.

One way to approach this is through work done on clarifying the notion of individuality [56]. As Griesemer notes this is a fraught concept [57], “I think we learn more about the concepts, claims, and consequences of individuality in the sciences by supposing it is a theoretical term that can mean different, even conflicting, things in different theoretical contexts, such as in different scientific specialties and disciplines with different research questions, projects, and phenomena to consider, rather than a logical or metaphysical term that that should shave a single, best, all-purpose fixed meaning” p.137. One useful perspective on individuality in the context of community ecology is Roberta Millstein’s articulation of “Land Community” as envisioned by Aldo Leopold [58, 59].

A land community combines aspects of community ecology and ecosystem ecology that emphasizes energy flow through the biotic and abiotic components of the system and the relationships among organisms and their abiotic context. Both of these are important in thinking about these as autonomous systems and provide a framework that provides a practical conception of boundaries. Millstein defines a Leopoldean land community as consisting of, “populations of different species interacting with each other and with their abiotic environment; these survival-relevant interactions often produce a flow of energy and materials between biotic components and between biotic components and abiotic components (and vice versa).”

She then articulates a notion of boundaries for these systems, “Land community boundaries for well-bounded systems are where discontinuities or steep gradients in the flow of material and energy coincide with discontinuities or steep gradients in species interactions. Land Community boundaries for open systems are at a minimum delineated by the smaller of the two types of discontinuities or steep gradients, including the more extensive interactions or matter/energy flows if and only if those interactions or matter/energy flows are stronger or larger than those of the smaller area.”

Lastly, she sets up three criteria for a land community to be considered an individual that we will use as a basis to explore such as autonomous systems. They are, (1) spatially and temporally restricted; (2) integrated by causal interactions which produce a shard fate; (3) are temporally bracketed with a beginning, and an end; and (4) are continuous in time allowing for change. We read the fourth as considering these as process-based. These set the stage for what further is required to consider these as autonomous, and in all that follows, by community we mean a Land community as described by Millstein.

Nunes-Neto, et al. [54] make a compelling argument that ecological communities can be seen as autonomous systems because of the functional components that structure most ecosystems. In particular, they look at the function of the components of biodiversity using what is termed the ‘organizational approach’ (OA).

Mossio, Saborido, and Moreno [60], formally define biological function as, “A trait T has a function in the organization O of a system S if and only if:

C1: T contributes to the maintenance of the organization O of S;
C2: T is produced and maintained under some constraints exerted by O;
C3: S is organizationally differentiated. [60] p. 828

This is contrasted with the Etiological Approach (EA), which uses evolutionary history to define function as doing that which it was selected to do in an adaptionist sense. OA then considers only what the function does in the current ecosystem, not as it is embedded in its historical or evolutionary framing. This, drawing on the work of Moreno and Mossio [53] on constraint closure, leads to an ecological definition of function, by Nunes et al. “An ecological function is a precise (differentiated) effect of a given constraining action on the flow of matter and energy (process) performed by a given item of biodiversity, in an ecosystem closure of constraints” [54].(p. 131)

This provides a handle into seeing land communities as autonomous systems, as maintained by constraint closure. The constraints are modulations induced by the components of biodiversity acting on the flow of matter and energy through the system, such that they feed back into the maintenance of the ecosystem. It is important to point out that the claim of its being an autonomous system is not the same as the claim that they are individuals in the biological sense, although they do seem to exhibit weak individuality [51, 54].

Dussault and Bouchard [50] add a significant consideration to the above account, developing a focus on particular kinds of functions that they call Persistence Enhancing Functions (PEP). PEPs are a class of functions that allow the currently structured ecosystem to persist. The focus is not upon these functions in an adaptive sense, such that their historical grounding has been necessarily worked out. Rather it is future-directed such that it attends to those extant functions that contributed to current stability-defined as the ability to return to equilibrium after a disturbance. They define a PEP function as, “The ¡¡PEP¿¿ function of x in an ecosystem E is to F if, and only if, x is capable of doing F and x’s capacity to F contributes to E’s propensity to persist.” [50](p. 8)

PEPs then allow us to explore the question, what features provide for the ability of this ecosystem, envisioned as a Leopold land community, to survive disturbances to equilibrium. It pulls away from questions about how particular assemblages come to be constructed and looks for general characteristics of ecosystem functioning and construction that are based upon the roles that species play in the current community.

These two proposals (OA and PEP) offer possible connections with the evolutionary constraints we develop below regarding ecological communities, and that influence stability and structure and that allow them to be considered autonomous systems.

### Vellend’s Set of Processes as Constraints

By combining the above ideas about how functions contribute to ecosystem structure and autonomy with Vellend’s proposal on evolutionary processes, we can see a possible theoretical space open with which to unite a view of evolutionary processes as constraints that stabilize the system. It might reasonably be argued that the evolutionary processes Vellend identifies can be thought to constrain and structure the ‘flow’ of species through an ecosystem. Because the constraints that Moreno and Mossio discuss are not limited to the kinds of systems typically discussed in the flow of matter and energy, e.g., chemical or cellular systems, and other micro-processes, any set of constraints on a given process, including the flow of species through an ecosystem, would seem to be a candidate for the emergence of autonomy in the sense they prefigure.

Selection constrains the success of a species by affecting its ability to compete in its current abiotic and biotic situation; providing fitness conditions on a species’ success or failure in surviving in an area. Species drift constrains the population genetic structure and what variation is available to a community’s species, and how stable the presence of a species is in a bounded region—including how it drifts in adjusting or modifying its current niche conditions. Speciation, in addition to the effects of selection, constrains how many new species emerge in an area due to genetic factors, and the interplay between trait differences and the biological fitness of those differences. Likewise, dispersal constrains species flow by influencing a combination of a species’ ability to spread beyond its original location, for instance in the sense that geneticist Sewall Wright proposed in his three-step shifting balance theory, i.e., (1) Initial Establishment; (2) Population Growth; and (3) Species dispersal of individuals to new areas [61]. This forms a hypothesis about how constraints influence ecosystem function. Constrains on species flow through the system may set up such communities as autonomous systems through constraint closure analogous to the way Moreno and Mossio [53] envisioned how autonomous systems are maintained for a period of time and, in particular, frame a set of PEP functions that influence stability and other aspects in community dynamics. There may be other possible constraints on species flow. For example, things like boundaries over which species cannot cross may affect which species enter into a particular community; the spatial patch dynamics of particular landscapes such that metapopulation structure creates a constraint on species flow; the frequency of disturbance; and mutualistic relationships between species within an ecosystem, which affects survival probabilities of cooperating species.

### A Model of Constraint Closure in Autonomous Ecological Communities

The mycorrhizal networks that form mutualistic relationships with plants may be a good target system to explore ideas about constraints and stability in communities [62, 63]. Soil mycorrhizal fungi form symbiotic relationships with tree roots and establish a chemical communication system among a diversity of ecosystem constituents [64]. These networks can enhance plant growth by increasing nutrient availability by facilitating their release from rocky substrate and helping break down organic carbon sources [65]. They also may help improve soil quality by modifying heavy metals and other inorganic pollutants [66, 67]. They have been shown to also affect interactions among competing plants, invasive species, and can have a large impact on community composition [68–71]. This is because these fungal communities can indirectly change the relationships among competing plants by providing resources to a broader spectrum of plants, thereby increasing above ground niche breadth [72].

Below we use processes typically found in mycorrhizal networks as a model system to test how the combination of Vellend’s suggestions about using a base set of processes as the building blocks for developing a generalized theory of community interactions in ecological systems. By using the ideas of constraint closure, we hope to provide a theoretical proof of concept for some of the patterns found in such communities and offer plausible interpretations for such patterns. The following is a generalized model. We are not trying to target a specific mycorrhizal ecosystem found in a field study reported in the scientific literature, but rather keeping an eye toward the kind of interactions found in the broad class of forest/mycorrhizal systems. This model system would lend itself to exploring nested mutualistic structures, but such nesting is not included in this paper [44].To put the following in terms of a model mapping schema provided by Weisberg (2013) [73] the model below is a generalized, computational model, using a minimalist idealization of key processes thought to potentially function in ecological communities, including Vellend’s [15] proposed set of high-level processes for conceptually framing ecological communities, i.e., selection, species drift, speciation, and dispersal. In addition, the model considers the kinds of constraints thought to capture the dynamics proposed by Moreno and Mossio [53]. These two ideas should allow community stability as recognized by signature patterns to emerge. The fidelity criteria we aim for will be achieved if such patterns are present and suggest that these processes are sufficient to confer known emergent attributes on these model systems. We make no claim that this model will actually represent any specific target system, but will give in principle insight into places for further testable hypotheses about how these ecosystems function.

To explore this possibility, we have developed an agent-based model that explores the potential of using these ideas to gesture to the possibility that the ideas of constraint closure can be a way to structure general insights into theoretical ecological community ecology.

## Results and Discussion

The role of the chosen constraints was clear from the simulation results. It took the shape of both inducing stability and creating conditions for a more dynamic community with increases in species turnover through time. In fig 2 through fig 6 key ecological variables showed overall stability in the landscape structure when plotted against the number of constraints. For example, fig 2a shows that with an increasing number of constraints, the overall fitness, and the variance (fig 3) in the fitness was decreasing on average across the landscape. Implying, by Fisher’s fundamental theorem, that there was a lowering of evolvability across the landscape. Likewise, in fig 4a and fig 5a, both species evenness and species richness were reduced. These measures all indicate that the number of constraints, regardless of which constraint considered, simplify the systems and create a more stable community. In addition to the count of constraints, the type of constraint played an important role in determining outcomes. Random forest models using the individual outcomes as dependent variables and the constraints as independent variables provide information about variable importance (a good description of the use and evaluation of random forests can be found in [74]). Constraint importance was determined by calculating the mean standard error (MSE) for each model after randomly permuting one of the constraints, and then calculating the percentage change from the unpermuted model (e.g. comparing the model’s MSE before and after shuffling the values for fitness competition). High values of importance indicate that a given constraint is necessary for maintaining model accuracy, while low values indicate that a constraint does not influence model error. Panel B in Fig 2, Fig 4, Fig 5, and Fig 6 shows the importance of each constraint for the four main simulation outcomes. Random forests show the overall importance of individual constraints, but they do not show the interactive effects of combinations of different constraints. To explore these interactions, panel C in Fig 2, Fig 4, Fig 5, and Fig 6 shows the average value of each simulation outcome for every combination of constraint.

**Fig 2.**
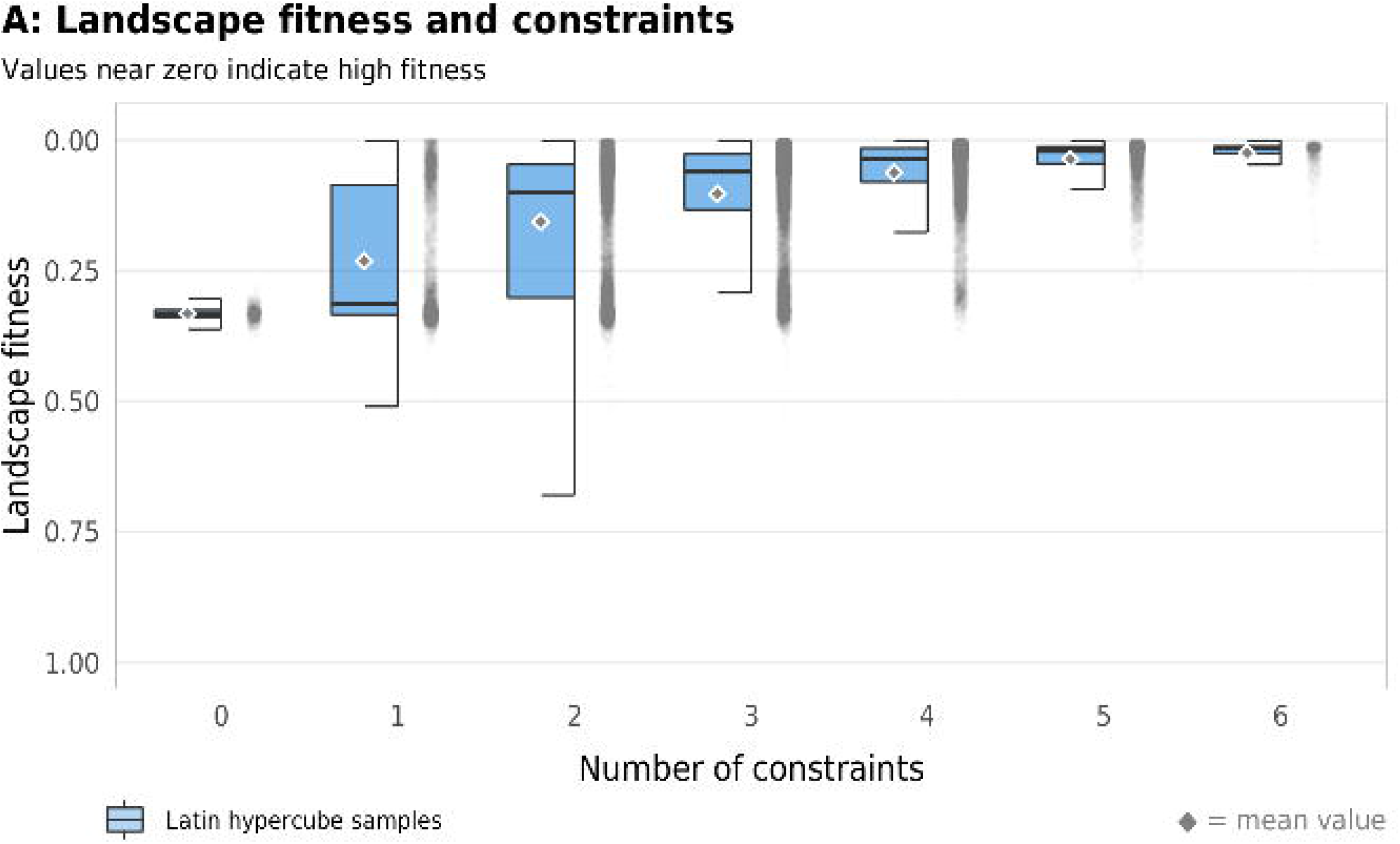
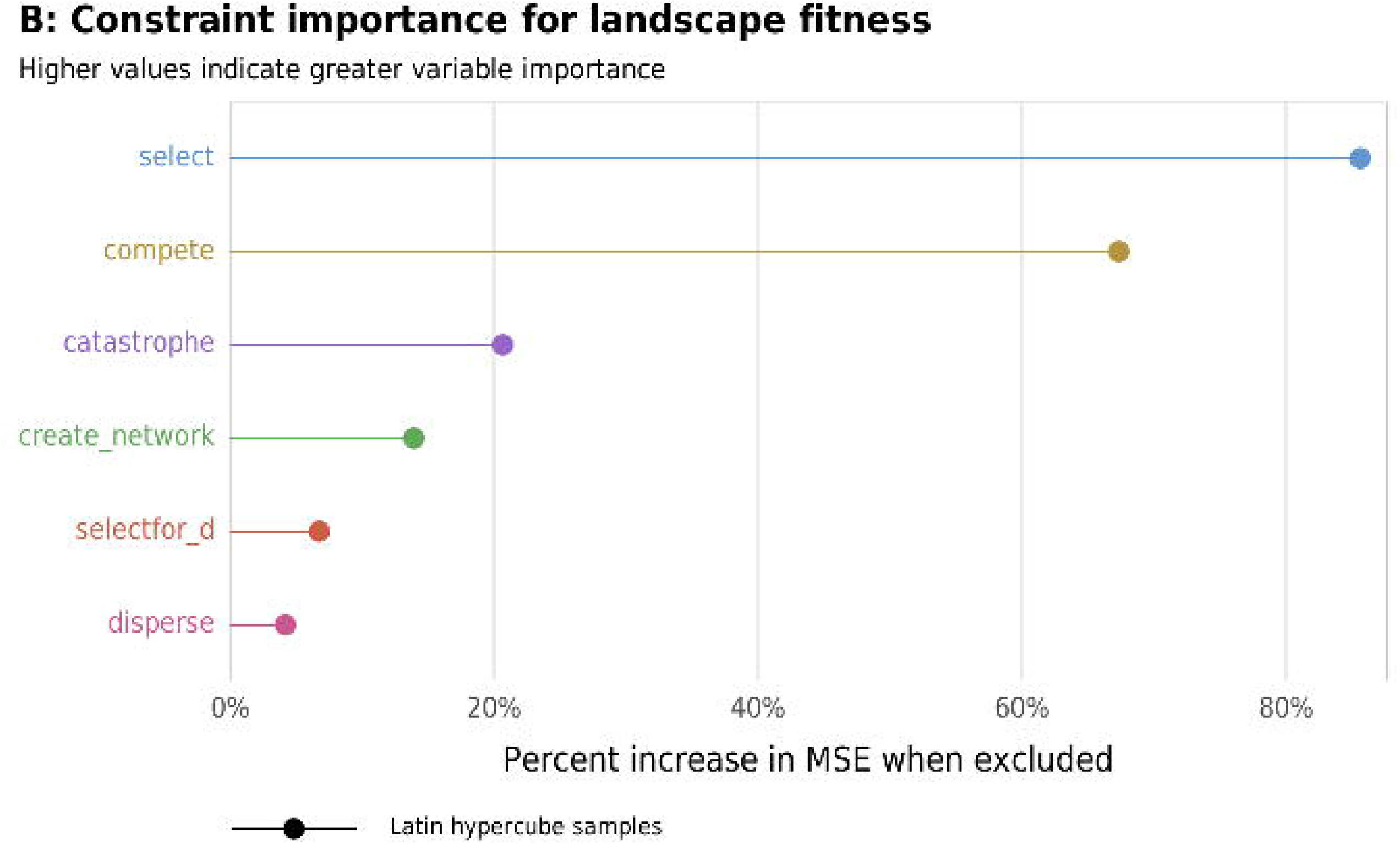
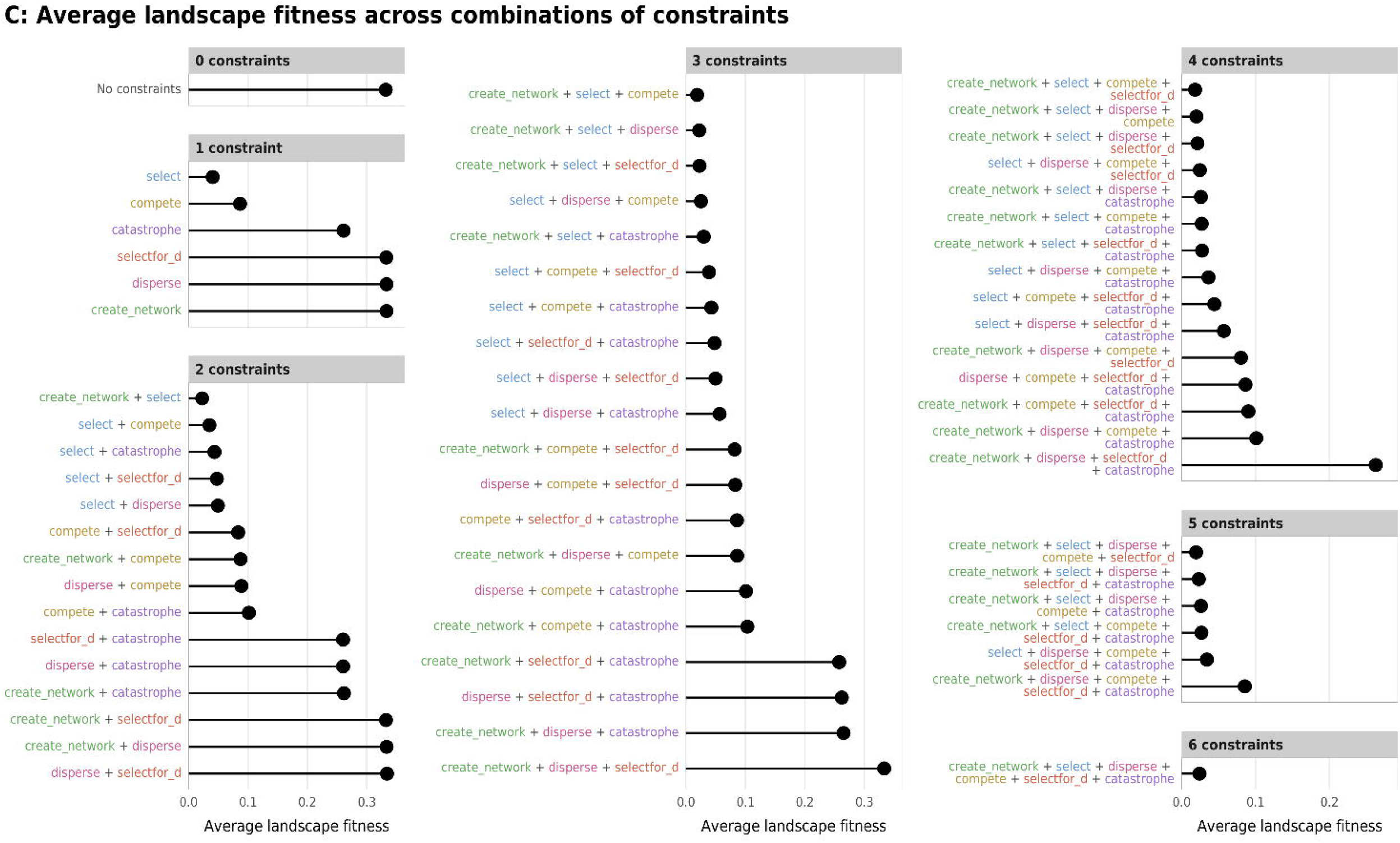
Effect of the number of constraints on landscape fitness. (A) Boxplots showing landscape fitness across different numbers of constraints; (B) Percent increase in mean standard error in a random forest model when each constraint is removed; (C) Average landscape fitness across all possible combinations of constraints.

**Fig 3.**
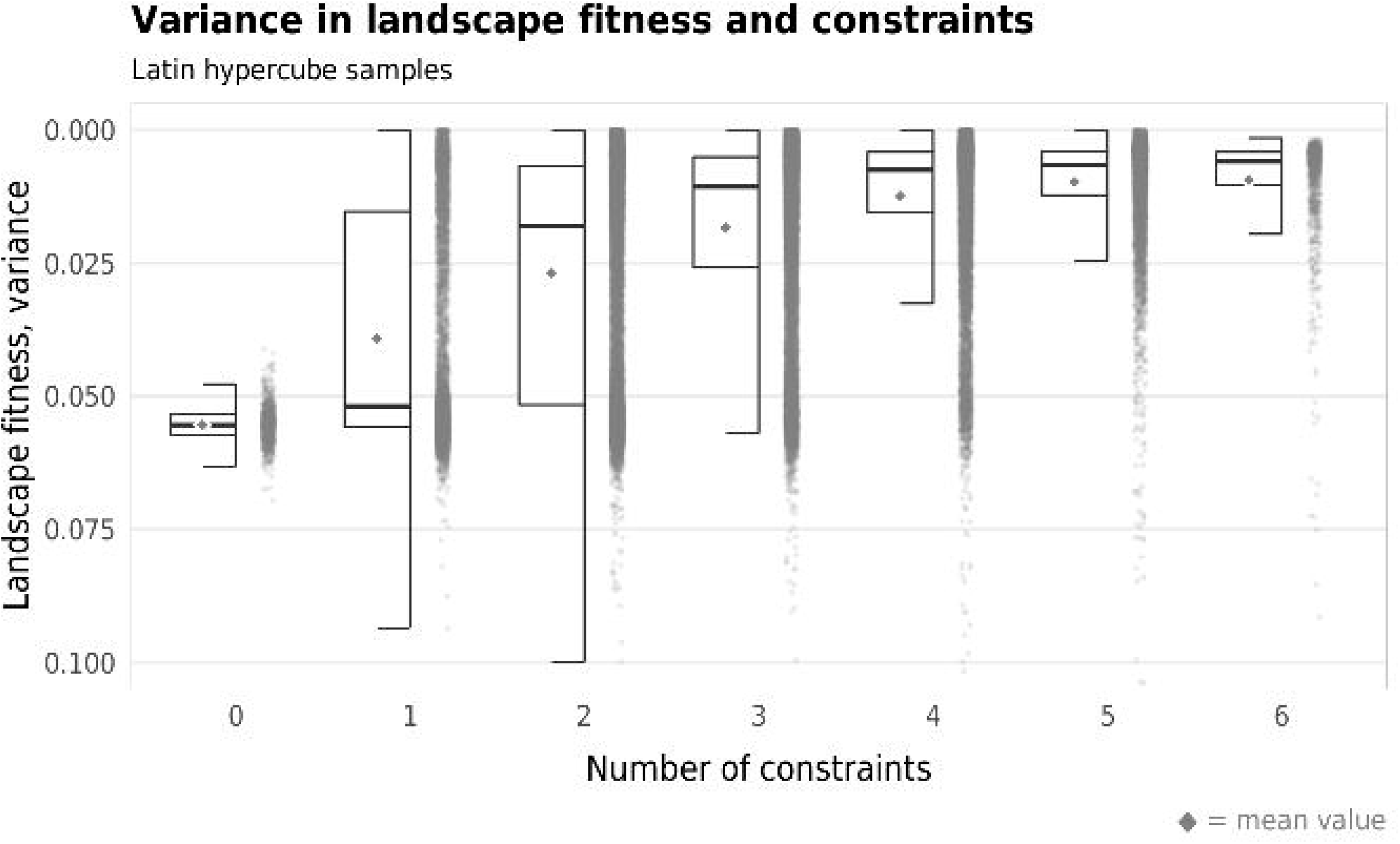
Effect of the number of constraints on variance of landscape fitness. Boxplots showing variance of landscape fitness across different numbers of constraints.

**Fig 4.**
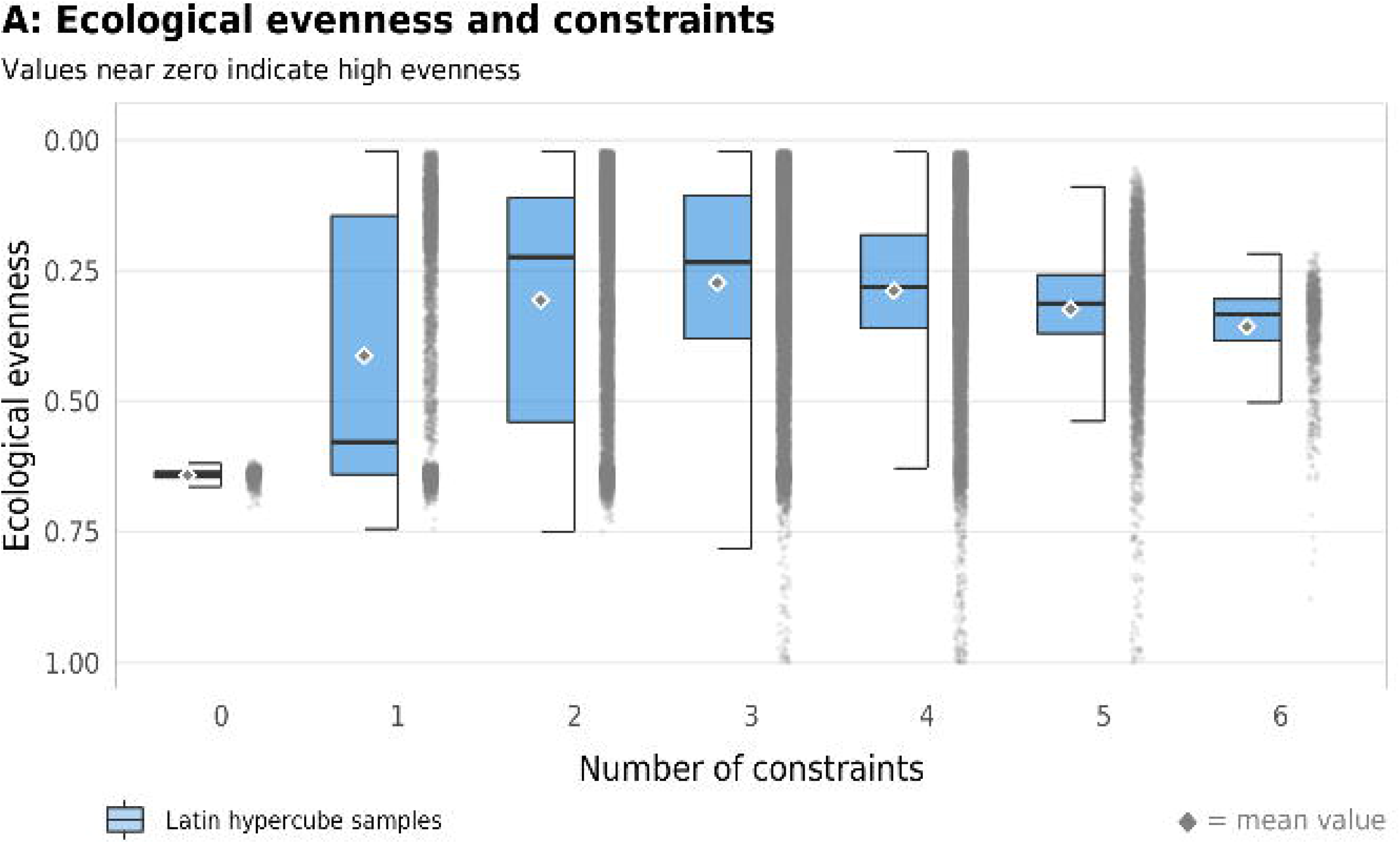
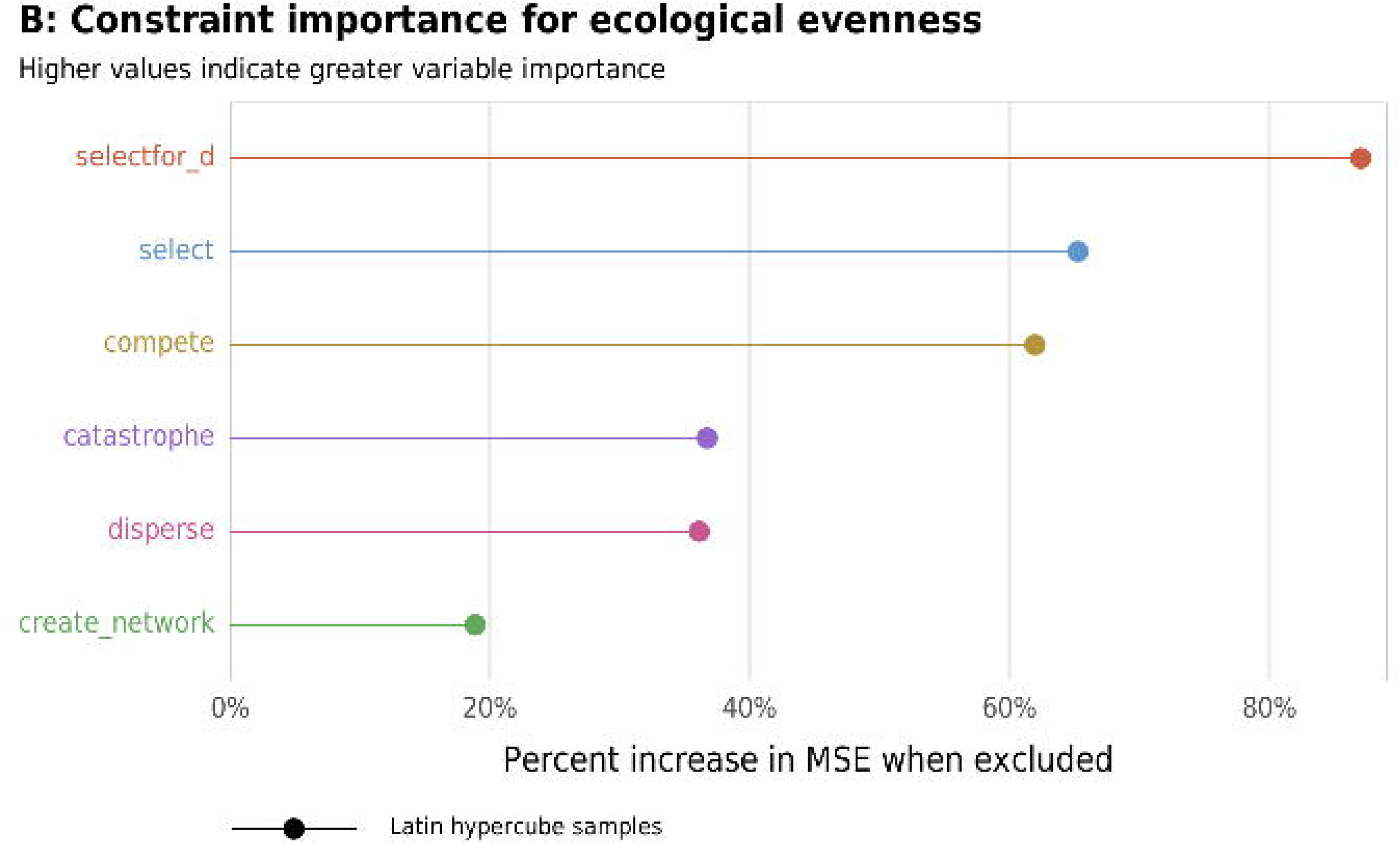
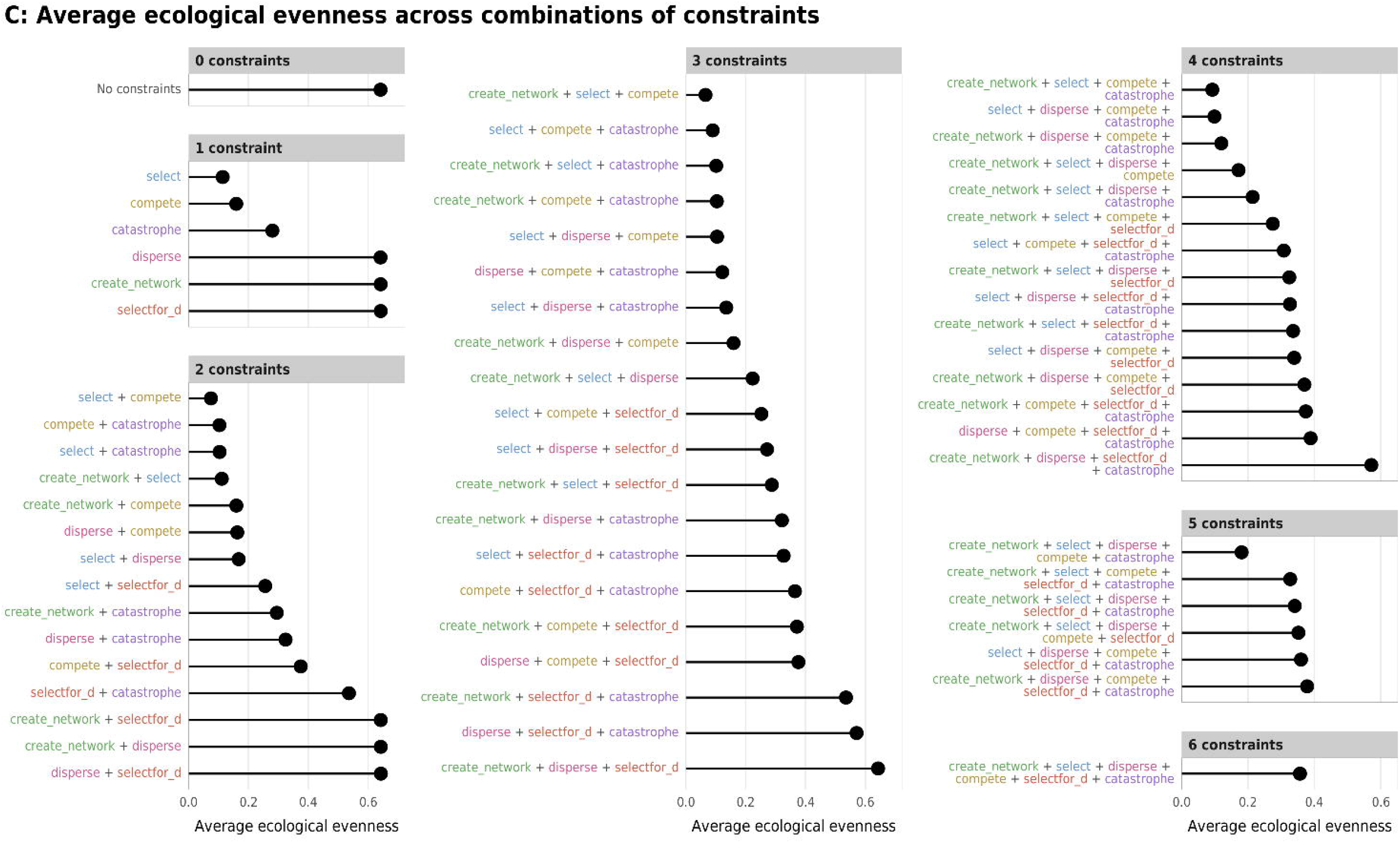
Effect of the number of constraints on ecological evenness. (A) Boxplots showing ecological evenness across different numbers of constraints; (B) Percent increase in mean standard error in a random forest model when each constraint is removed; (C) Average ecological evenness across all possible combinations of constraints.

**Fig 5.**
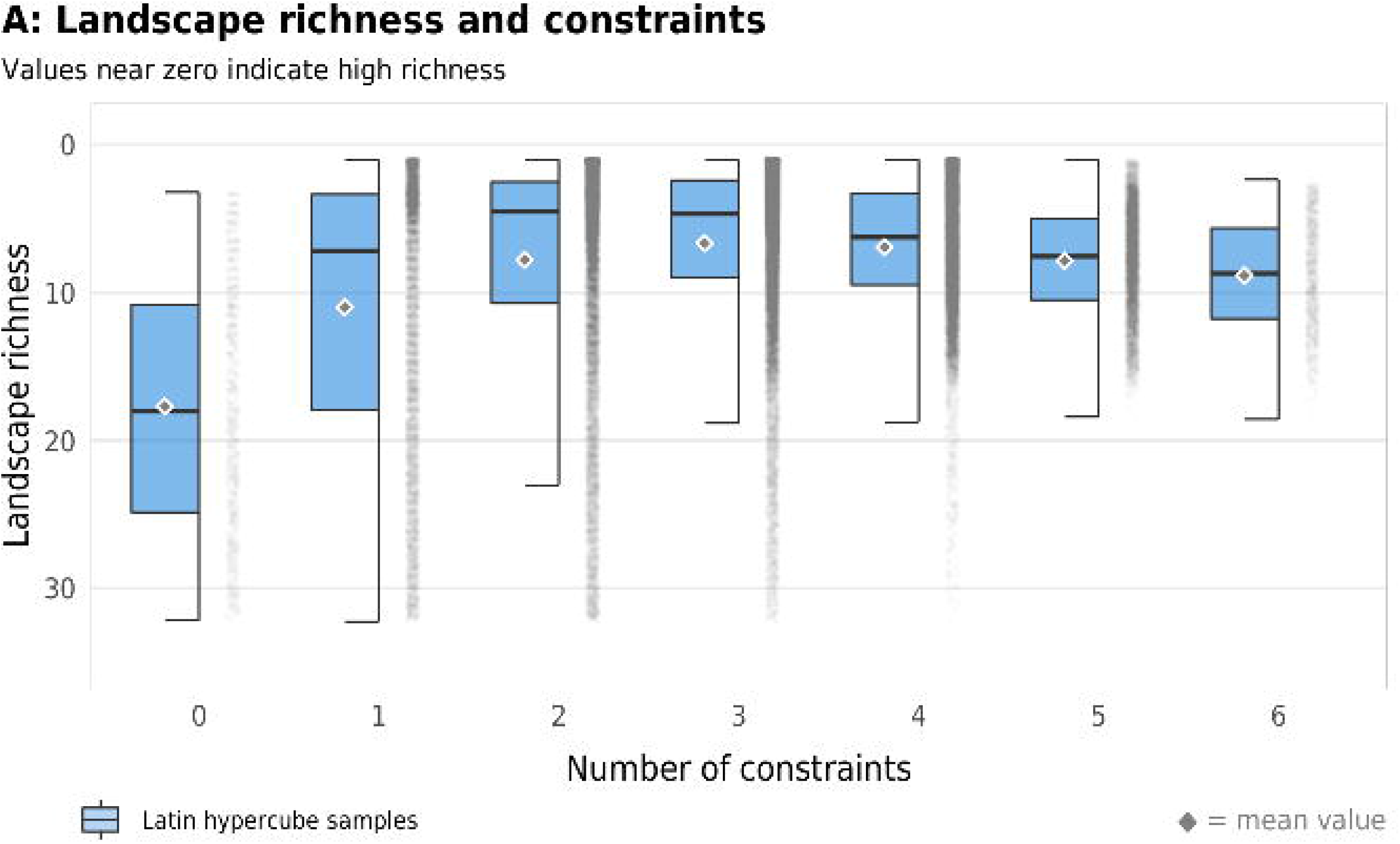
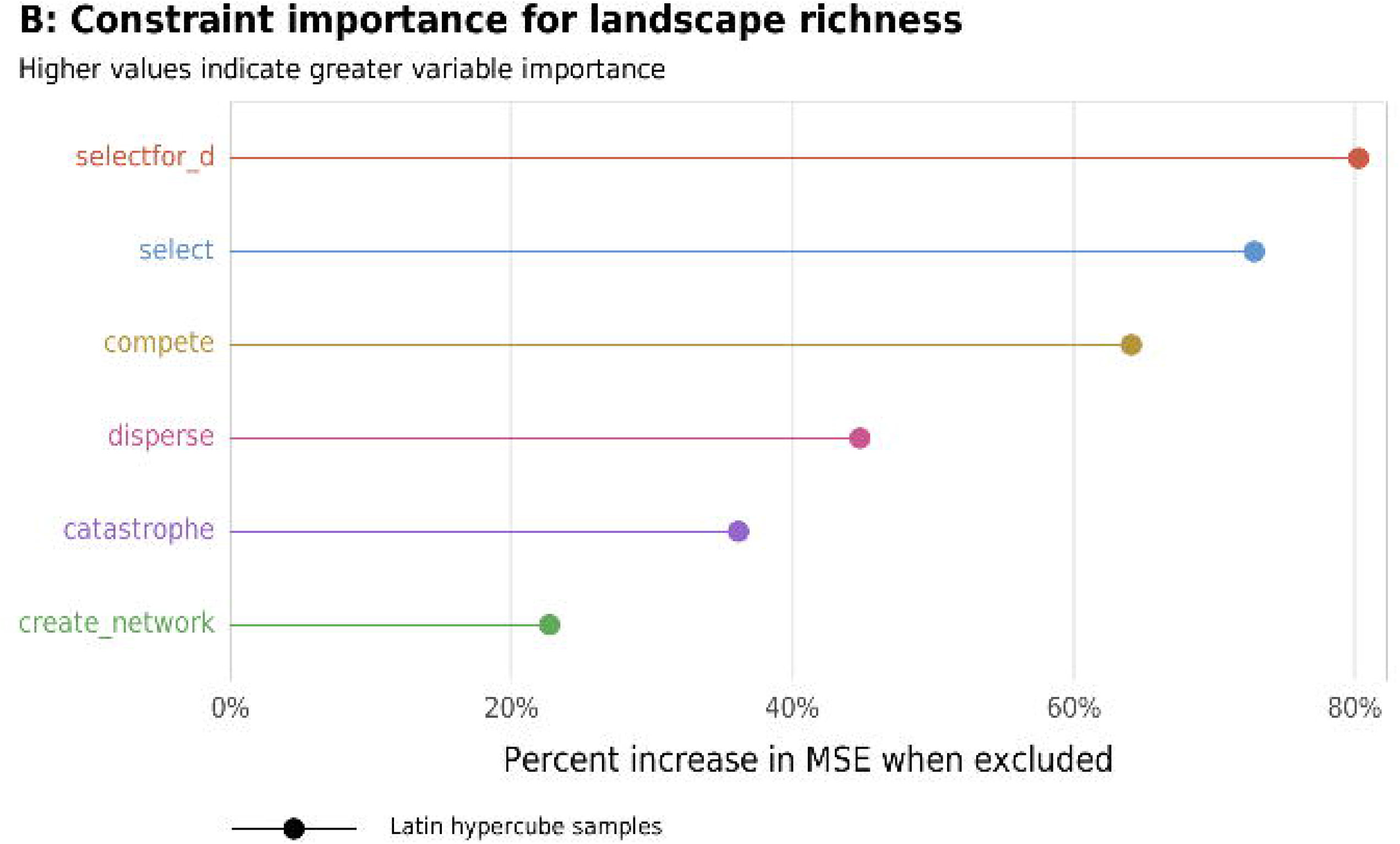
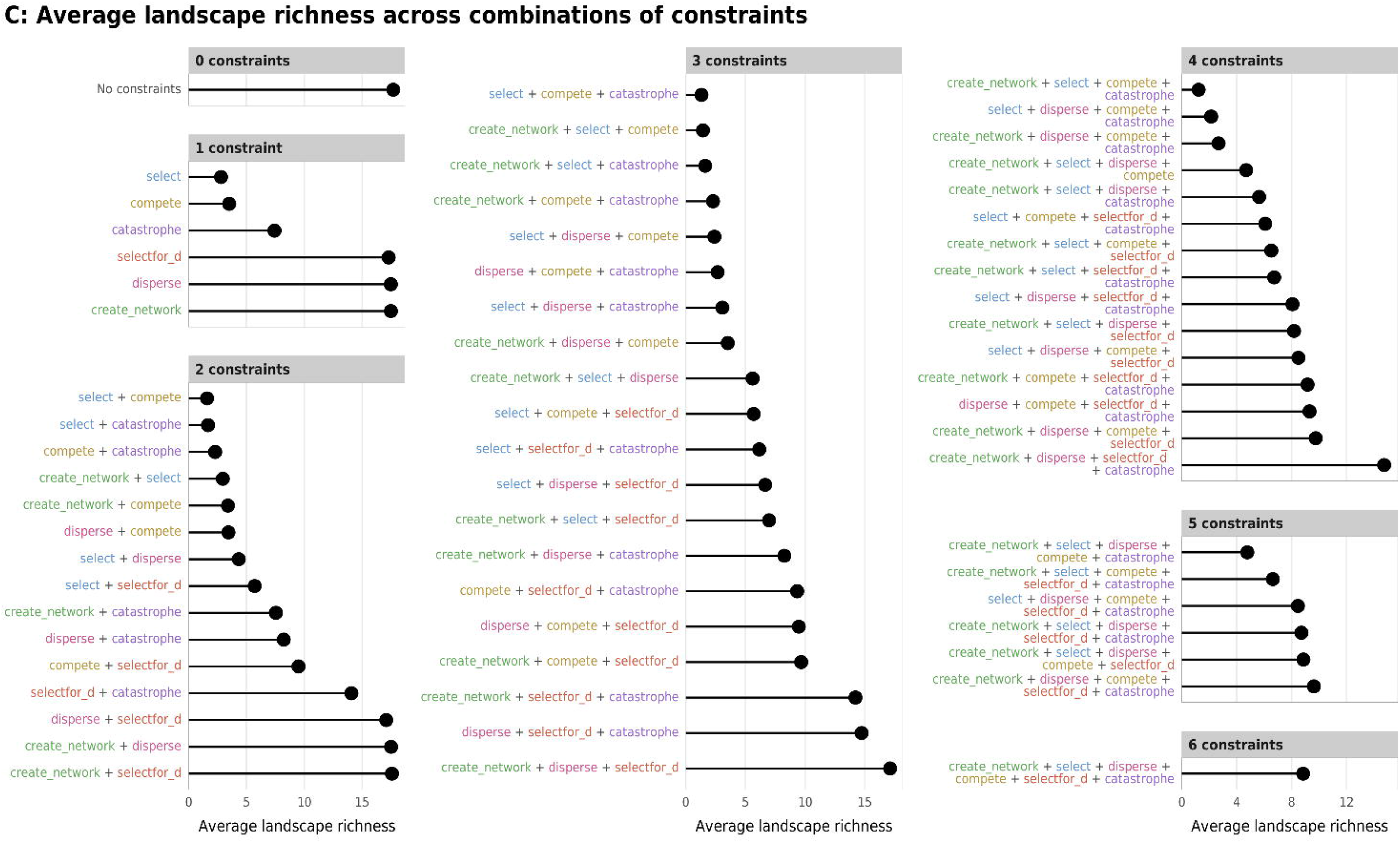
Effect of the number of constraints on species richness. (A) Boxplots showing species richness across different numbers of constraints; (B) Percent increase in mean standard error in a random forest model when each constraint is removed; (C) Average species richness across all possible combinations of constraints.

**Fig 6.**
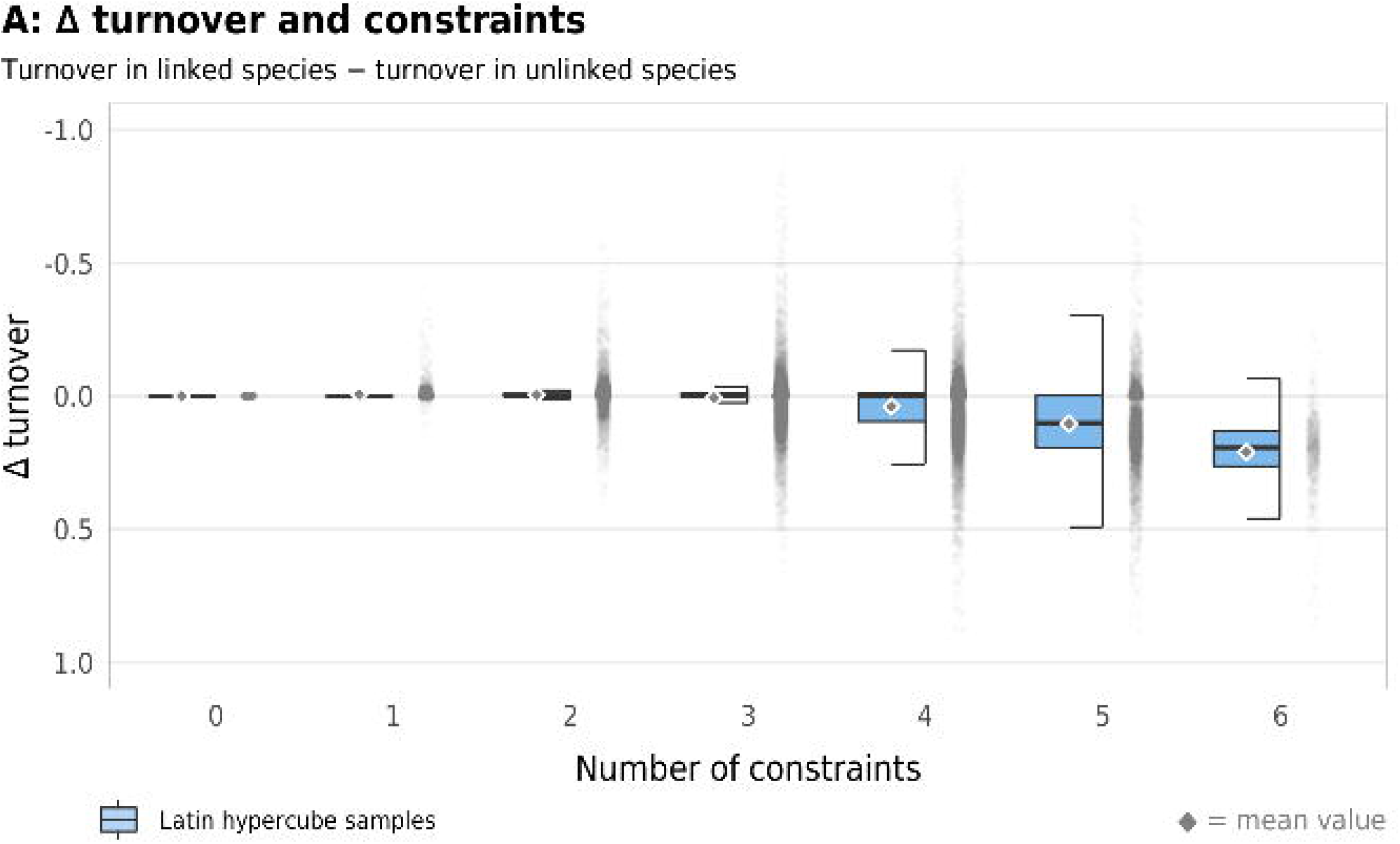
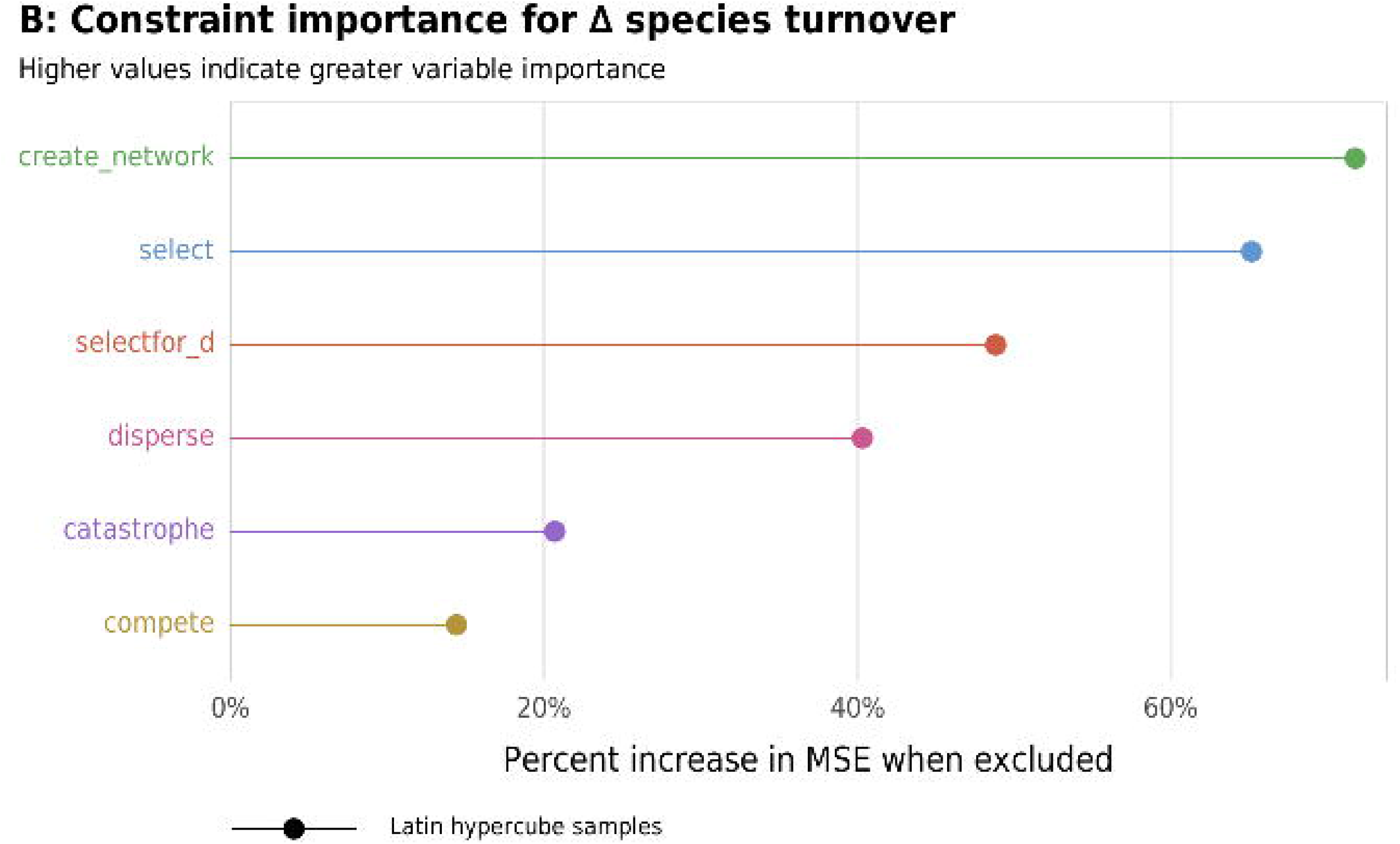
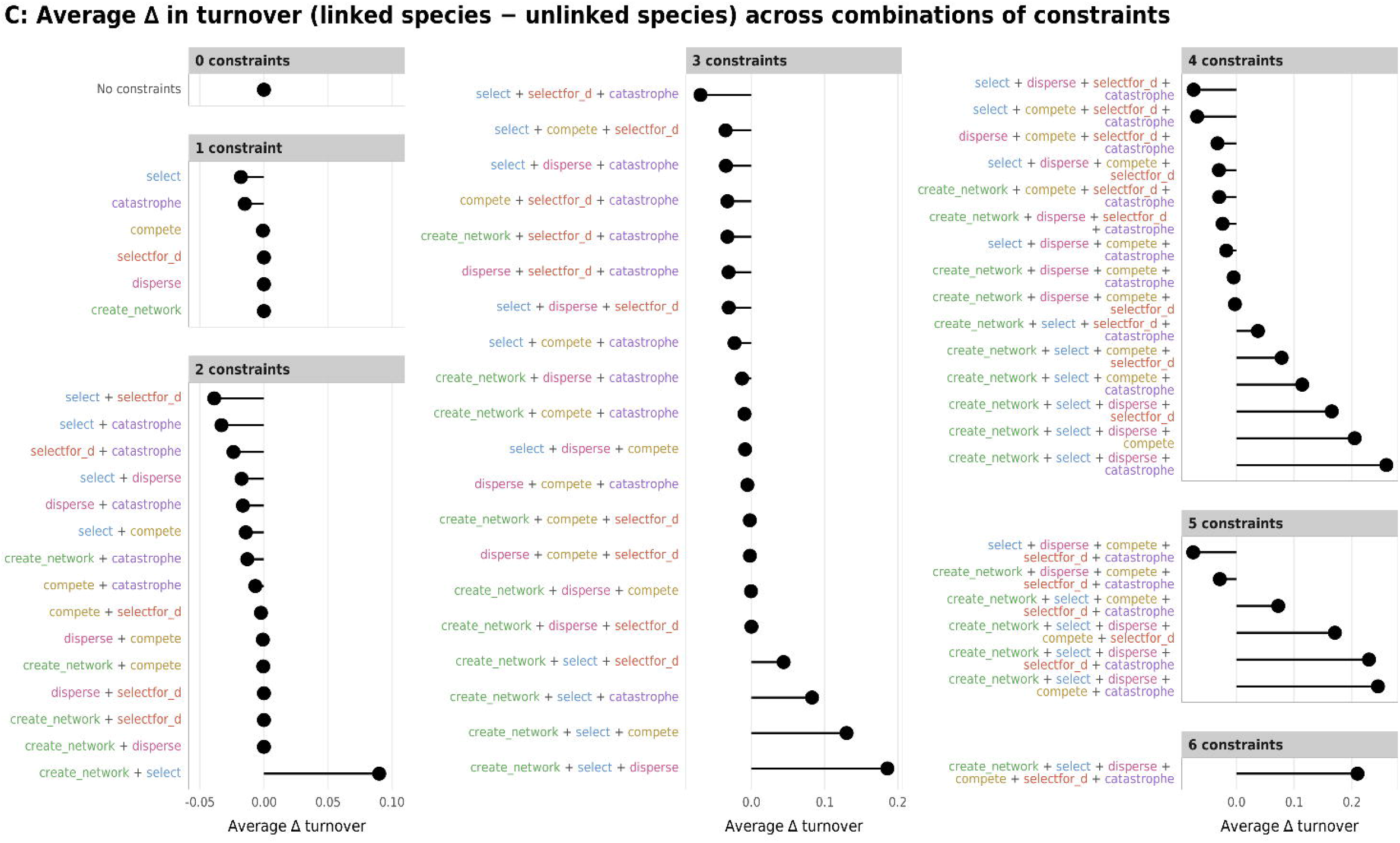
Effect of the number of constraints on the difference between turnover in linked and unlinked species. (A) Boxplots showing difference between turnover in linked and unlinked species across different numbers of constraints; (B) Percent increase in mean standard error in a random forest model when each constraint is removed; (C) Average difference across all possible combinations of constraints.

## Conclusions

While the argument that these evolutionary constraints needs more work in terms of both theortical and modeling work, this work points to the possibility that these constraints may effect measures of ecosystem stability and maintenance.

It might be objected that this type of analogy might not hold in the way that systems such as biological cells are maintained through constraint closure through specific mechanisms that constrain the flow of energy in material systems. Stretching the notion of constraint closure to something as nebulous as multivalent as evolutionary ‘forces’ on individual species and their formation and extinction should be approached with caution. However, we believe it has enough promise to be worth turning over to theorists and modelers who could formalize the structural arguments to demonstrate whether the analogy holds. If these evolutionary forces constrain species in the way we suggest, then the structures of community ecology such as stability and long-term persistence should emerge as constant patterns in such systems allowing for a research program for community ecology that would be able to discover general features of these systems that maintain their continuance through time.

The emergence of particular pattern formation suggests empirical work that may lend evidentiary support as to whether these evolutionary forces form PEP functions in the right way to meet the above criteria defined by Dussault and Bouchard (2016) [50]. Especially, though the use modeling (e.g., agent-based modeling), which provides the structure that would allow the exploration of constraint closure and its effect on ecologically relevant pattern formation, community stability, and turnover.

## Methods and Model Description

### A word about modeling in theory construction

Representing an ecological community in a model presents several challenges. Modeling such systems has a rich history, and three types of models have dominated: Analytic, those largely represented by mathematical systems of equations (called analytic even if computation must be used to solve the equation using numerical methods); Simulation or computational models, those represented by algorithms instantiating procedures, rules, and/or manipulating abstract objects or data structures representing individuals or agents; or statistical models that look for relationships among data objects using probability theory to structure inferences and predictions from the data. All three of these are used regularly in community ecology and serve different purposes.

In all models what is being represented varies over the uses the developer wants target with her models. For example, the target systems my be a specific community with specified entities and for which the modeler hopes to make specific claims or make targeted predictions about this particular system. On the other end of the spectrum, the target might be a generalized ideation of a hypothesized structure. For example in early in the history of population genetics, long before the nature of DNA was worked out, much progress was made modeling the properties that genes must necessarily possess to do the work proposed for what it meant to be a gene.

Given the complexity of ecological systems, simulation seems the method best equipped to handle their complexity. Simulation is best viewed as a hermeneutical practice [75].

Pattern recognition has played a pivotal role in ecology almost since its inception [76], but especially since Levin’s influential paper laying out the important role in ecosystems of spatial patterns and scale [77]. Since that time, ecological researchers have focused on an number of issues related to pattern formation both temporal and spatial scales, including, scales of interaction in both in trophic relationships; modularity of ecosystems function; patterns of diversity; assemblage rules; and other influences that are dependent on differential scales of influence and interaction [78]. This leads to a number modeling problems orbiting questions about how to best represent these systems, especially in regard to the use of null models, or in trying to sort out assemblage rules, which empirically have proven to be complex and difficult to accomplish adequately [4], making modeling even more challenging. Especially if, as argued above, these have emergent properties and patterns. Modeling emergent phenomena require that we take into account the diachronic aspects of the system that allow the emergence and modeling of individual interactions. This is most easily done using agent-based models. They may be an important way of drawing out the connection between the patterns we find in nature and those generated by our idealized models [79], p. 150.

Grimm and Railsback [80] have offered a rationale and strategy for recognizing and modeling patterns in complex ecological systems, which they call ‘pattern-oriented modeling’ (POM). This lends itself well to agent-based models and for exploring the complexity inherent in ecological communities and identifying the right levels and processes to include in the model. They suggest the following set of practices be used after empirically identifying the patterns:

”(i) determine what scales, entities, variables and processes the model needs, (ii) test and select submodels to represent key low-level processes such as adaptive behavior, and (iii) find useful parameter values during calibration.” p. 298.

For example, consider the problem of exploring Vellend’s proposal in light of the work on constraint closure in mycorrhizal networks as a model system to explore these questions.

We develop an agent-based model designed to explore how evolutionary processes form closed constraints that provide long-term stability in ecological communities.

## ODD

We will use the “Overview, Design Concepts, and Details” (ODD) protocol recommended by Grimm et. al. 2010 [81] to describe the model. This protocol has proven to be an important way to describe computer simulation studies and in particular agent-based models such as this one. This protocol facilitates understanding of how the model has been designed, what assumptions the authors are making, and how various components of these kinds of models, such as spatial scale, time, stochasticity, and other choices for structuring the model are implemented. The model is written in NetLogo 6.0.2 an Agent-Based Programming Environment [82].

### Purpose

The following model has been designed to provide in-principal arguments for how ecological communities maintain stability even in the presence of stochastic turnover. Using Vellend’s [15] proposed set of high-level processes for conceptually framing ecological communities, i.e., selection, species drift, speciation, and dispersal, we hypothesize that these provide constraint closure in the way proposed by Moreno and Mossio [53] that contribute to biological autonomy. Moreover, we hypothesize specifically that these will allow stable ecological signatures to arise and be maintained through constraint closure. In this way, we seek to exemplify the possibility of creating some measure of generalizability in ecosystem community structure.

### Entities, state variables, and scales

Table 2 gives a list of the parameters in the model. The model is structured on a spatial toroidal grid composed of *N*x*N* patches that form clusters of suitable and unsuitable areas. procedure is based upon O’Sullivan and Perry [83].

**Table 2.** Table of Principal Parameters Used in the Model.

| Parameter | Name (role) | Range | Default Value |
| --- | --- | --- | --- |
| $T_{max}$ | Maximum number of time steps | $\in \{1, 20, 000\}$ | 500 |
| $S_c$ | Species Selection (constraint) | Binary Switch | On |
| $D_c$ | Species Dispersal (constraint) | Binary Switch | On |
| $C_c$ | Species Competition (constraint) | Binary Switch | On |
| $\Omega_c$ | Speciation (constraint) | Binary Switch | On |
| $P_c$ | Perturbation (constraint) | Binary Switch | On |
| $\Psi$ | Proportion Species which form mutualisms | $\in \{0, 1\}$ | 0.5 |
| $\nu$ | variance in environmental needs (used in fitness) | $\in \{0, 1\}$ | 0.05 |
| $\Phi$ | Frequency of Perturbation | $\in \{0, T_{max}\}$ | 500 |
| $\Sigma$ | Proportion of species wiped out in perturbation | $\in \{0, 1\}$ | 0.2 |
| $\lambda$ | Landscape Density Factor | $\in \{0, 1\}$ | 0.25 |
| $\Lambda$ | Patch Distribution Factor | $\in \{0, 20\}$ | 5 |
| $\xi$ | Maximum Number of Species per Patch | $\in \{1, 50\}$ | 10 |
| $\kappa$ | Number of Mutualisms | $\in \{1, \xi_{max}\}$ | 5 |

Table notes: All parameters are coded either integers or random variables

The unsuitable habitat patches form a barrier or impediment to migration. Each habitat patch represents a packed set of niches that can hold *M* species. Each patch contains an environmental niche score, a variable representing a habitat suitability index chosen from a uniform random distribution between 0 and 1. Each species has a similar niche score, which represents a one-dimensional niche measure of the species needs. This corresponds to the patch environmental niche score, which in turn is a measure of what needs that habitat will provide. A species’ fitness will be defined as the absolute difference between a species’ niche score and its patch niche score.

A subset of the species in in the model are able to form a mutualistic relationship with other species and an undirected network is established between *k* <= *M* species. The fitness of a linked species, differs from unlinked species and is calculated as a given function of linked species in a given patch, e.g., the maximum or average fitness of the linked species.

Each species also plays a functional role in a patch. This is specified by *sp_ν_* in which *ν* ∈ {1… *sp_max_*} and *sp_max_* is the maximum number of functional roles in each patch’s niches. The functional roles also determine a “link designation” which picks out which species can form links, i.e., *sp_numlink_*, designates which species can have links, and which cannot, such that *sp_ν_* ≤ *numlink* links can be formed among other species that can also form links.

The specific constraints thought to allow the creation of stable communities are (a) Selection; (b) Dispersal; (c) mutualism (with any eye to mycorrhizal symbiotic relationships); (d) Spatial Competition; and (e) Fitness Competition between linked and unlinked species. In addition, there are two assumed constraints that are structural parts of the model and are not tested for effect in this stage of the model analysis but may be explored at a later time. (a) niche packing, i.e., a finite niche space; and (b) spatial structure limiting dispersal among species.

### Process overview and scheduling

The overall flow of the model is given in fig 7.

**Fig 7.**
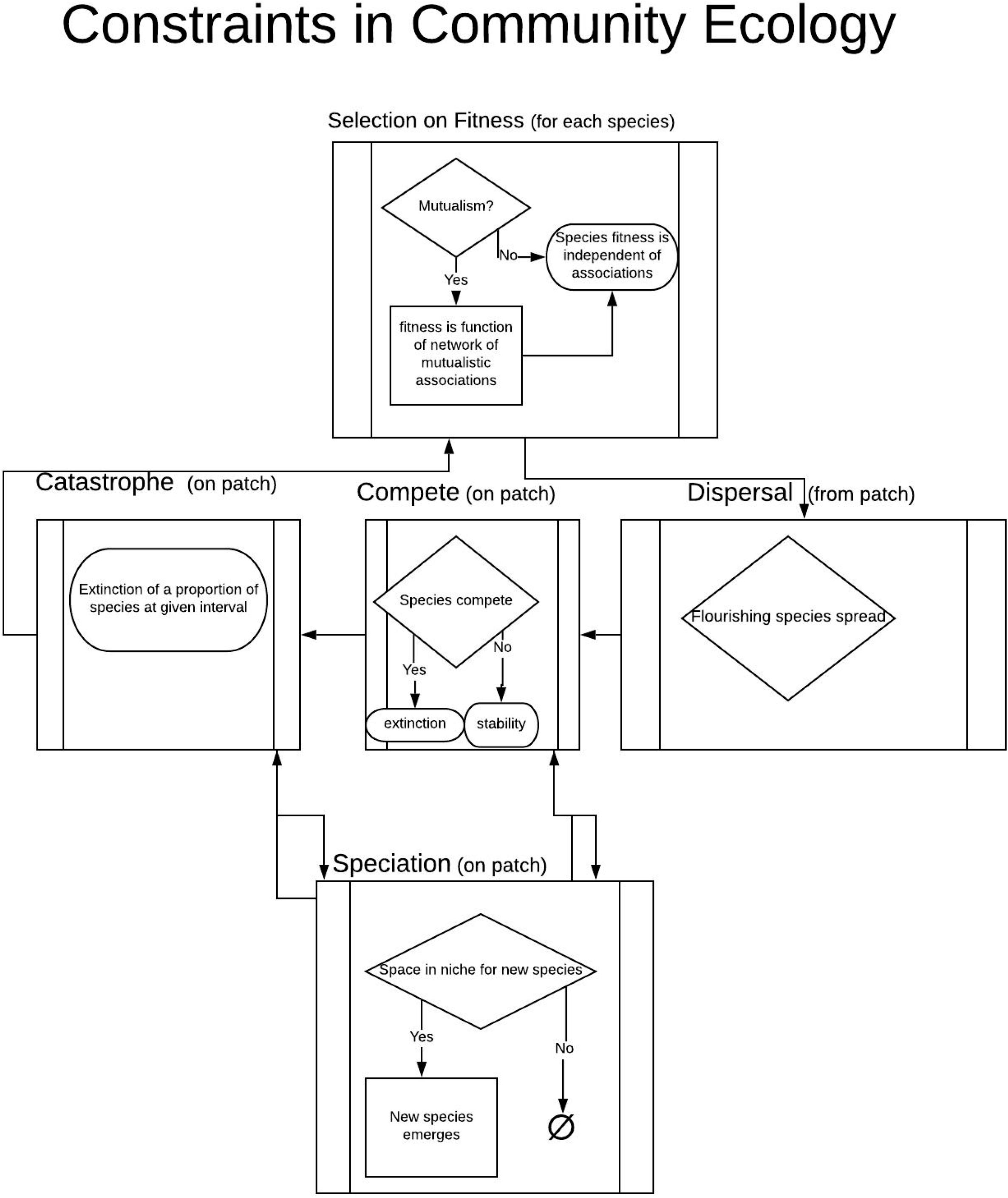
Process Flow Within Model. The agent-based model used in this paper cycles through the evolutionary components of Vellend’s proposed evolutionary structures in ecological systems (described above). These structures are used as constraints to create semi-autonomous structures. The flow chart shows the order of events and possible flow paths possible in the model.

After establishing the grid (details below in initialization section), *M* species are created in each patch *i*. A given number *K*, are able to form mutualistic links, the rest, *M* – *K*, are not. Each time step, *τ*, represents the mean time a species is likely to exist in a community before patch extinction or species evolution. At time, *τ*_0_, *k* mutualistic links are established within the patch for the initial population of species in each patch where *k* = *K_ρ_*. *K* is the maximum number of links that can be established, and *ρ* is a random variable that gives the proportion of potential links that do get established for a given *K*. The following processes procedures are given in the order they are executed:

- Select: Fitness is calculated based on the absolute distance of a given species’ niche score to the patch’s environmental niche score. For linked species the fitness is the highest fitness among those species with which it is linked. The species with the lowest fitness on each patch is eliminated.
- Dispersal: The species with the highest fitness spreads to one of its neighbors if there exists an opening, i.e., one of its neighbor patches has fewer than *M* species. If it is a species that can form links in its new patch, it will do so.
- Fitness competition: One of the linkable species and one of the unlinkable species is randomly selected within each patch. The one with the higher fitness reproduces, the one with the lower fitness goes extinct in the patch. The new species will have a niche score with the same mean fitness as its parent’s fitness value, with variance *σ*.
- Species drift: Whenever there are *G* <= *M* openings in the niches within a patch, *G* new species are created from the *G* species with the highest fitnesses. The new species will have a niche score with the same mean fitness as its parent’s fitness value, but with variance *σ*. There will also be the possibility the species drifting to a new functional role. The drift can occur across the link designator boundary *sp_numlink_*, i.e., species can go from a linkable species to unlinkable species and visa versa, but if it drifts such that *ν* < 1 or *ν* < *maxsp* it reflects back to the numlink boundary without changing its link designation, i.e., If *sp*_1_ where about to drift to the non-existent *sp*_0_, then *sp*_1_ ⇒ *sp*_*numlink*-1_ and likewise if *sp_maxsp_* was going to drift out of the range above *maxsp*, then *sp_maxsp_* ⇒ *sp*_*numlink*+1_ such that it remains in its current link designation.
- Catastrophes: At given intervals, *I* ∈ 1, *t_max_* a proportion, *ϕ*, of the *M* species in each patch can be randomly wiped out. These perturbations provide a test of stability in terms of how the patch recovers from its loss of species.

### Design Concepts

Emergence: We expect stability to emerge from certain combinations of constraints that form a closed loop of interaction and create constraint closure among the ecological processes described above.

- Objectives: We run the model under a number of different parameter combinations to understand how stability is structured.
- Interactions: Among species-agents in the model, interactions can be positive among those forming mutualistic associations, and negative among those that compete.
- Stochasticity: Stochasticity enters into the model in several places. (a) The initial patch structure of the landscape is randomly generated; (b) The proportion of species chosen to actually form links, *ρ* is a random variable; (c) the species that go extinct in the drift stage are randomly chosen with probability proportional to fitness; species with high fitness chosen to move, move randomly to one of their 8-Moore-neighborhood neighbors; (d) The variation of fitness around newly spectated species is a random variable *ν*; and initially the environmental variable and the species environmental needs variable are randomly chosen; and (e) Species eliminated in a catastrophe are randomly chosen.

### Variables of Interest

- *Turnover rate*: The turn over rate is the number of new species created at each time step. The way the model is formulated, new species arise when there is an opening in relation to the packed niche. This happens in competition between the linked and unlinked species, in selection events in which the least fit species is selected for elimination, and in perturbation events.
- *Evenness*: To capture patch evenness the Pielou’s evenness index was modified for algorithmic efficiency. Rather than the mathematical equation 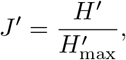, where H’ is Shannon’s Index, 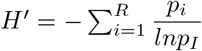 we modify it as follows. Because the number of species per patch is fixed, the algorithm used is computed by the number of unique species in a patch divided by the number of species in the patch. This has the range, 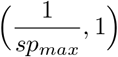 of where the lower range is found in the case where every species in the patch is unique, and the higher when there is only one species.
- *Landscape Fitness* Fitness is measured as the mean fitness among all species in the landscape.

### Model Structure and Testing

- Initialization: Initialization consists of a random landscape being produced using two variables that control the number of patches selected and the distribution of patches.
- Submodels: The only submodels are those described in the Process Overview section.
- Testing: The model Behavior Space in NetLogo was used to test the overall behavior of the model. A complete iteration set of the binary switches turning on and off the constraint submodels and for three values of the possible number of links per species and two values of different landscape types leading to 144 possible combinations which compromise the permutation set used to explore the constraint space, with *t_max_* = 500 timesteps for each sample from the model hyperspace. A uniform random distribution of the variables propform, varfitness, frqmod, and perwipe, were drawn from the 150 iterations of each of the possible constraint permutation set for each of the These were taken with replacement over the multidimensional parameter space. As White et al. (2014) note [84], these results should not be interpreted as a test of statistical significance of each of the parameter set as the likelihood values and statistical tests are easily influenced by the number of iterations used in the model testing. Therefore p-values were provided as a heuristic to choose the set of parameters that might be averaged or representative values chosen that do not influence how the constraints inform the results of the model.

### Statistical Analysis of Latin Hypersphere Runs

In the Latin Hypersphere run of the model with random non-constraint independent variables, the following analyses were used to reconstruct the effects of the constraints on the model landscape. For each of the constraints, random forests were generated to explore how different combinations and subsets of the chosen evolutionary constraints affected the ecological indices [85].

